# The mutational and phenotypic spectrum of *TUBA1A*-associated tubulinopathy

**DOI:** 10.1101/427948

**Authors:** Moritz Hebebrand, Ulrike Hüffmeier, Steffen Uebe, Arif B. Ekici, Cornelia Kraus, Mandy Krumbiegel, André Reis, Christian T. Thiel, Bernt Popp

## Abstract

**Background:** The *TUBA1A*-associated tubulinopathy is clinically heterogeneous with brain malformations, microcephaly, developmental delay and epilepsy being the main clinical features. It is an autosomal dominant disorder mostly caused by *de novo* variants in *TUBA1A*.

**Results:** In three individuals with developmental delay we identified heterozygous *de novo* missense variants in *TUBA1A* using exome sequencing. While the c.1307G>A, p.(Gly436Asp) variant was novel, the two variants c.518C>T, p.(Pro173Leu) and c.641G>A, p.(Arg214His) were previously described. We compared the variable phenotype observed in these individuals with a carefully conducted review of the current literature and identified 166 individuals, 146 born and 20 fetuses with a *TUBA1A* variant. In 107 cases with available clinical information we standardized the reported phenotypes according to the Human Phenotype Ontology. The most commonly reported features were developmental delay (98%), anomalies of the corpus callosum (96%), microcephaly (76%) and lissencephaly (70%), although reporting was incomplete in the different studies. We identified a total of 121 distinct variants, including 15 recurrent ones. Missense variants cluster in the C-terminal region around the most commonly affected amino acid position Arg402 (13.3%). In a three-dimensional protein modelling, 38.6% of all disease causing variants including those in the C-terminal region are predicted to affect binding of microtubule-associated proteins or motor proteins. Genotype-phenotype analysis for recurrent variants showed an overrepresentation of certain clinical features. However, individuals with these variants are often reported in the same publication.

**Conclusions:** With 166 individuals, we present the most comprehensive phenotypic and genotypic standardized synopsis for clinical interpretation of *TUBA1A* variants. Despite this considerable number, a detailed genotype-phenotype characterization is limited by large inter-study variability in reporting.

## INTRODUCTION

The superfamily of tubulin genes is composed of alpha-, beta-, gamma-, delta- and epsilon families. The alpha and beta families, consisting of at least 15 alpha and 21 beta-tubulin genes, respectively,^1^ encode tubulin proteins which form heterodimers as fundamental components of microtubules.^2^ Along with microtubule associated proteins (MAPs) and motor proteins on the external surface, tubulin proteins participate in substantial cellular processes of intracellular transport, cell division and neuronal migration.^3,4^

In recent years, an increasing number of tubulin genes were linked to a clinically heterogeneous group of disorders, the “tubulinopathies” (*TUBA1A*, MIM#602529; *TUBA8*, MIM#605742; *TUBB2A*, MIM#615101; *TUBB2B*, MIM#612850; *TUBB3*, MIM#602661; *TUBB*, MIM#191130; *TUBG1*, MIM#191135).^5-11^ Tubulinopathies are characterized by a broad spectrum of cortical and subcortical malformations and a variety of clinical features. Major cortical anomalies include lissencephaly, polymicrogyria or polymicrogyria-like cortical dysplasia and cortical gyral simplification. Subcortical anomalies affect the corpus callosum, the cerebellar vermis, the brainstem, the basal ganglia and the cerebellum. Further clinical features are microcephaly, developmental delay and epilepsy.^12,13^ To date, *TUBA1A* represents the main tubulinopathy gene and accounts for 4-5% of all lissencephaly cases.^14, 15^

Using exome analysis in three unrelated individuals with severe developmental delay we identified three heterozygous *de novo* missense variants in the *TUBA1A* gene. We extensively reviewed and systematically reanalyzed available public data to provide a standardized synopsis of described variants together with reported neuroradiological and clinical features of TUBA1A-associated tubulinopathy. We used this comprehensive information to perform a detailed analysis of the genotypic and phenotypic spectrum highlighting a possible genotype-phenotype relationship and probable bias in reporting.

## MATERIALS AND METHODS

### Clinical reports

**Individual i084n**: A 13-year-old boy was the second child of healthy non-consanguineous parents of European descent and was born at term after an uneventful pregnancy. At age six months, the boy presented with a complex focal seizure and a multifocal infantile cerebral seizure disorder was diagnosed. Anticonvulsive therapy with vigabatrin and valproate was started leading to an initial remission followed by refractory Blitz-Nick-Salaam-like seizures. Cerebral magnetic resonance imaging (MRI) revealed coarsened cerebral gyri, a hypoplastic rostrum of the corpus callosum, cerebellar vermis hypoplasia, moderately enlarged lateral ventricles and narrowed white matter. At age 4 years severe hypotonia, global developmental delay, spastic movement disorder, strabismus convergens and pyelectasia became apparent. On referral at age 13, he presented with a weight of 41.0 kg (10^th^ - 25^th^ centile), a height of 146.3 cm (3^rd^ centile) and an occipitofrontal circumference (OFC) of 52 cm (<3^rd^ centile). Minor facial features included a flat forehead, low set ears, epicanthic fold, upward slanting palpebral fissures, strabismus, narrow nasal bridge, broad nose tip, short philtrum and an everted lower lip. In addition, a clinodactyly V, a small forefoot and a sandal gap were noticed. Family history and previous genetic workup including chromosome and chromosomal microarray (CMA) were unremarkable.

**Individual i085n:** A 14.5-year-old boy was the second child of healthy non-consanguineous parents of European descent. He was born at term after an uneventful pregnancy [4170 g (85^th^ - 97^th^ centile), 55 cm (97^th^ centile), and OFC 34 cm (10^th^ - 25^th^ percentile)]. Motor and language milestones were delayed and cerebral MRI revealed mild frontal cortical anomalies, hypoplastic corpus callosum, basal ganglia dysgenesis, ventricular dilatation, accentuated lamina quadrigemina, massive chambered retrocerebellar arachnoidalcyst and a pineal gland cyst. The boy was last seen at age 11.5 years and presented with a weight of 87.6 kg (>97^th^ centile), height of 172 cm (97^th^ centile) and head circumference of 56 cm (25^th^ - 50^th^ percentile). His facial gestalt included a high forehead, large earlobes, epicanthic folds, hypertelorism, jaw deformity, cupid bow shaped upper lip with open mouth appearance, high arched palate and a gap between the upper incisors. Minor features were distally located thumbs, pointed fingers, hallux valgus, and a sandal gap. Family history and previous genetic testing including CMA were unremarkable.

**Individual i086n:** A 10-year-old girl was the first child of healthy non-consanguineous parents of European descent. She was born at term with parameters in the normal range [3050 g (15^th^ - 40^th^ centile), 51 cm (75^th^ - 90^th^ centile), OFC 34 cm (25^th^ - 50^th^ centile)]. At age six months, a generalized hypotonia and nystagmus were noted followed by developmental regression at 1 year 9 months. Myoclonic and later generalized seizures were treated with anticonvulsive therapy (topiramate, valproate). Cerebral MRI revealed a Dandy walker malformation, with cerebellar vermis hypoplasia, agenesis of corpus callosum and ventricular dilatation. Further features were fused basal ganglia, and unilateral optic nerve hypoplasia. At last physical examination (age 9 years 9 months), she presented with a height of 125 cm (<3^rd^ centile) and head circumference of 50.5 cm (10^th^ - 25^th^ percentile). Severe language delay with absent speech and muscular hypotonia with spasticity of the legs were noted. Minor facial features included large ears, hypertelorism, broad flat nasal-bridge, high arched palate, wide spaced teeth, thin lips, short neck, smooth philtrum and a simian crease. Additional features included proximal located thumbs, pointed fingers, brachymesophalangia V and clinodactyly V. Previous testing including CMA was unremarkable. The identification of the *TUBA1A* variant in this girl was part of a previous publication without detailed clinical description (reported as ID S_006).^16^

### Exome Sequencing

Informed written consent was obtained for all participants. The study was approved by the Ethical Committee of the Medical Faculty of the Friedrich-Alexander-Universität Erlangen-Nürnberg. DNA from peripheral blood lymphocytes was extracted using standard methods. Exome sequencing was performed after SureSelect v5 (i085n, i086n) and v6 (i084n) targeted capturing on HiSeq 2500 for i084n and i085n (Trio analysis^17^) and i086n (Exome Pool-Seq^16^). After mapping of sequence reads to the GRCh37/hg19 reference genome and variant calling using standard methods for the trio analysis^17^ or as described by Popp et al. for the exome Pool-Seq^16^, variants in coding regions including splice sites were selected based on population frequency (gnomAD) and computational prediction scores, e.g. CADD score^18^. Variants were confirmed, and segregation tested by Sanger sequencing.

### Review of reported *TUBA1A* cases from literature and databases

We identified 112 articles, published between 01/2007 and 06/2018, from PubMed applying the search term “TUBA1A”. Of these, 28 provided clinical reports and were thus included in this study. All available clinical data was standardized in accordance to terms of the Human Phenotype Ontology (HPO).^19^ In contrast to a previously established classification combining cortical and subcortical features like “classic lissencephaly”, “lissencephaly with cerebellar hypoplasia”, “lissencephaly with agenesis of the corpus callosum” and “centrally predominant pachygyria”^12,20^, we analyzed the features independently. If only the classification was mentioned we used the independent underlying features where HPO terms were available (e.g. “microlissencephaly”: microcephaly HP:0000252 + agyria HP:0031882). Nevertheless, we kept composite terms typically used together in the literature such as “agyria-pachygyria” (HP:0031882, HP:0001302) if they affected the same brain structure. Data assessment comprised 11 neuroradiological features, including anomalies of cortical gyration, corpus callosum, brainstem, basal ganglia, internal capsule, cerebellum, cerebellar vermis, hippocampus, ventricular dilatation, 4^th^ ventricle dilatation, grey matter heterotopia, and other radiological findings. Clinical features included congenital microcephaly, microcephaly, developmental delay, epilepsy, neuro-ophthalmological findings including strabismus and nystagmus, other neurological symptoms including spasticity and muscular hypotonia, and additional features (HPO terms shown in Table 1, 2 and File S1).

**Table 1.** Neuroradiological features of *TUBA1A*-associated tubulinopathy

| Clinical information | Born n (%) | Fetuses n (%) | i084n | i085n | i086n | Total n (%) |
| --- | --- | --- | --- | --- | --- | --- |
| Number of reported cases | **84 (100)** | **20 (100)** |  |  |  | **107 (100)** |
| Sex | 33f/36m/15n | 7f/12m/1n | m | m | w | 41f/50m/16n |
| **Abnormality of the Corpus Callosum (HP:0001273)** | **79/83 (95.2)** | **20/20 (100.0)** | + | + | + | **102/106 (96.2)** |
| agenesis (HP:0001274) | 15/83 (18.1) | 16/20 (80.0) |  |  | + | 32/106 (30.2) |
| partial agenesis (HP:0001338) | 14/83 (16.9) | 1/20 (5.0) |  |  |  | 15/106 (14.2) |
| dysplastic (HP:0006989) | 14/83 (16.9) | 3/20 (15.0) |  |  |  | 17/106 (16.0) |
| hypoplasia (HP:0002079) | 31/83 (37.5) | 0/20 (0.0) | + | + |  | 33/106 (31.1) |
| partial agenesis, hypoplastic (HP:0001338, HP:0002079) | 5/83 (6.0) | 0/20 (0.0) |  |  |  | 5/106 (4.7) |
| normal | 4/83 (4.8) | 0/20 (0.0) |  |  |  | 4/106 (4.7) |
| no information available | 1/84 (1.2) | 0/20 (0.0) |  |  |  | 1/107 (0.9) |
| **Abnormal cortical gyration (HP:0002536)** | **74/75 (98.7)** | **19/19 (100.0)** | + | + |  | **95/96 (99.0)** |
| lissencephaly (HP:0006818) | 50/75 (66.7) | 17/19 (89.5) |  |  |  | 67/96 (70.0) |
| agyria (HP:0031882) | 12/75 (16.0) | 15/19 (78.9) |  |  |  | 27/96 (28.1) |
| agyria-pachygyria (HP:0031882, HP:0001302) | 15/75 (20.0) | 1/19 (5.3) |  |  |  | 16/96 (16.7) |
| pachygyria (HP:0001302) | 23/75 (30.7) | 1/19 (5.3) |  |  |  | 24/96 (25.0) |
| polymicrogyria (HP:0002126) | 16/75 (21.3) | 2/19 (10.5) |  |  |  | 18/96 (18.8) |
| perisylvian-polymicrogyria (HP:0012650) | 10/75 (13.3) | 0/20 (0.0) |  |  |  | 10/96 (10.4) |
| cortical gyral simplification (HP:0009879) | 5/75 (6.7) | 0/20 (0.0) |  |  |  | 5/96 (5.2) |
| unspecific | 3/75 (4.0) | 0/20 (0.0) | + | + |  | 5/96 (5.2) |
| normal | 1/75 (1.3) | 0/20 (0.0) |  |  |  | 1/96 (1.0) |
| no information available | 9/84 (10.7) | 1/20 (5.0) |  |  | + | 11/107 (10.3) |
| **Abnormality of the cerebellar vermis (HP:0002334)** | **58/62 (93.5)** | **18/18 (100.0)** | + |  | + | **78/83 (94.0)** |
| hypoplasia (HP:0001320) | 42/62 (67.7) | 12/18 (66.7) | + |  | + | 56/83 (67.5) |
| dysgenesis (HP:0002195) | 16/62 (25.8) | 6/18 (33.3) |  |  |  | 22/83 (26.5) |
| normal | 4/62 (6.5) | 0/20 (0.0) |  | + |  | 5/83 (6.0) |
| no information available | 22/84 (26.2) | 2/20 (10.0) |  |  |  | 24/107 (22.4) |
| **Abnormality of the basal ganglia (HP:0002134)** | **48/48 (100.0)** | **8/9 (88.9)** |  | + | + | **58/59 (98.3)** |
| dysgenesis (HP:0025102) | 48/48 (100.0) | 8/9 (88.9) |  | + | + | 58/59 (98.3) |
| normal | 0/48 (0.0) | 1/9 (11.1) |  |  |  | 1/59 (1.7) |
| no information available | 36/84 (42.9) | 11/20 (55.0) | + |  |  | 48/107 (44.9) |
| **Abnormality of the brainstem (HP:0002363)** | **39/47 (83.0)** | **18/18 (100.0)** |  |  |  | **57/65 (87.7)** |
| hypoplasia (HP:0002365) | 24/47 (51.1) | 8/18 (44.4) |  |  |  | 32/65 (49.2) |
| pons hypoplasia (HP:0012110) | 6/47 (12.8) | 10/18 (55.6) |  |  |  | 16/65 (24.6) |
| dysplasia (HP:0002508) | 9/47 (19.1) | 0/20 (0.0) |  |  |  | 9/65 (13.8) |
| normal | 8/47 (17.0) | 0/20 (0.0) |  |  |  | 8/65 (12.3) |
| no information available | 37/84 (44.0) | 2/20 (10.0) | + | + | + | 42/107 (39.3) |
| **Ventricular dilatation (HP:0002119)** | **40/40 (100.0)** | **6/6 (100.0)** | + | + | + | **49/49 (100.0)** |
| fourth ventricle dilatation (HP:0002198) | 18/40 (45.0) | 1/6 (16.7) |  |  | + | 20/49 (40.8) |
| no information available | 44/84 (52.4) | 14/20 (70.0) |  |  |  | 58/107 (54.2) |
| **Abnormality of the cerebellum (HP:0001317)** | **22/32 (68.8)** | **16/17 (94.1)** |  |  |  | **38/49 (77.6)** |
| dysplasia (HP:0007033) | 4/32 (12.5) | 6/17 (35.3) |  |  |  | 10/49 (20.4) |
| hypoplasia (HP:0001321) | 16/32 (50.0) | 10/17 (58.8) |  |  |  | 26/49 (53.1) |
| agenesis (HP:0012642) | 1/32 (3.1) | 0/20 (0.0) |  |  |  | 1/49 (2.0) |
| normal | 10/32 (31.3) | 1/17 (5.9) |  |  |  | 11/49 (22.4) |
| no information available | 52/84 (61.9) | 3/20 (15.0) | + | + | + | 58/107 (54.2) |
| **Abnormal morphology hippocampus (HP:0025100)** | **24/29 (82.8)** | **5/8 (62.5)** |  |  |  | **29/37 (78.4)** |
| hypoplasia (HP:0025517) | 6/29 (20.7) | 3/8 (37.5) |  |  |  | 9/37 (24.3) |
| dysgenesis (HP:0025101) | 18/29 (62.1) | 2/8 (25.0) |  |  |  | 20/37 (54.1) |
| normal | 5/29 (17.2) | 3/8 (37.5) |  |  |  | 8/37 (21.6) |
| no information available | 55/84 (65.5) | 12/20 (60.0) | + | + | + | 70/107 (65.4) |
| **Abnormality of the internal capsule (HP:0012502)** | **24/25 (96.0)** | **1/19 (100.0)** |  |  |  | **25/26 (96.2)** |
| anterior limb thinned or absent | 13/25 (52.0) | 0/20 (0.0) |  |  |  | 13/26 (50.0) |
| normal | 1/25 (4.0) | 0/20 (0.0) |  |  |  | 1/26 (3.8) |
| no information available | 59/84 (70.2) | 19/20 (95.0) | + | + | + | 81/107 (75.7) |
| **Grey matter heterotopia (HP:0002281)** | **11/13 (84.6)** | **14/15 (93.3)** |  |  |  | **25/28 (89.3)** |
| olivary | 5/13 (38.5) | 6/15 (40.0) |  |  |  | 11/28 (39.3) |
| absent | 2/13 (15.4) | 1/15 (6.7) |  |  |  | 3/28 (10.7) |
| no information available | 71/84 (84.5) | 5/20 (25.0) | + | + | + | 79/107 (73.8) |
| **Other radiological features** | **9/9 (100.0)** | **8/8 (100.0)** | + | + | + | **20/20 (100.0)** |
| abnormal morphology of the olfactory bulb (HP:0040327) | 2/9 (22.2) | 6/8 (75.0) |  |  |  | 8/20 (40.0) |
| no information available | 75/84 (89.3) | 12/20 (60.0) |  |  |  | 87/107 (81.3) |

**Table 2.** Clinical features of *TUBA1A-associated* tubulinopathy

| Clinical information | Born n (%) | Fetuses n (%) | i084n | i085n | i086n | Total n (%) |
| --- | --- | --- | --- | --- | --- | --- |
| Number of reported cases | 84 (100) | 20 (100) |  |  |  |  |
| Sex | 33f/36m/15n | 7f/12m/1n | m | m | w |  |
| **Microcephaly (HP:0000252)** | **46/52 (88.5)** | **10/20 (50.0)** | + |  |  | **57/75 (76.0)** |
| normal | 6/52 (11.5) | 10/20 (50.0) |  | + | + | 18/75 (24.0) |
| no information available | 32/84 (38.1) | 0/20 (0.0) |  |  |  | 32/107 (29.9) |
| **Congenital microcephaly (HP:0011451)** | **25/36 (69.4)** | n/a |  |  |  | **25/36 (69.4)** |
| normal | 11/36 (30.6) | n/a |  |  |  | 11/36 (30.6) |
| no information available | 48/84 (57.1) | n/a | + | + | + | 51/87 (58.6) |
| **Developmental delay (HP:0001263)** | **49/50 (98.0)** | n/a | + | + | + | **52/53 (98.1)** |
| normal | 1/50 (2.0) | n/a |  |  |  | 1/87 (1.1) |
| no information available | 34/84 (40.5) | n/a |  |  |  | 34/87 (39.1) |
| **Other neurological symptoms** | **36/37 (97.3)** | n/a | + | + | + | **39/40 (97.5)** |
| spasticity (HP:0001257) | 19/37 (51.4) | n/a |  |  |  | 19/40 (47.5) |
| muscular hypotonia (HP:0001252) | 9/37 (24.3) | n/a |  | + |  | 10/40 (25.0) |
| spasticity and muscular hypotonia (HP:0001257, HP:0001252) | 4/37 (10.8) | n/a | + |  | + | 6/40 (15.0) |
| other | 4/37 (10.8) | n/a |  |  |  | 4/40 (10.0) |
| normal | 1/37 (2.7) | n/a |  |  |  | 1/40 (2.5) |
| no information available | 47/84 (56.0) | n/a |  |  |  | 47/87 (54.0) |
| **Epilepsy (HP:0001250)** | **35/49 (71.4)** | n/a | + |  | + | **37/52 (71.2)** |
| generalized tonic-clonic seizures (HP:0002069) | 18/49 (36.7) | n/a |  |  | + | 19/52 (36.5) |
| infantile spasms (HP:0012469) | 4/49 (8.2) | n/a |  |  |  | 4/52 (7.7) |
| generalized tonic-clonic seizures and infantile spasms (HP:0002069, HP:0012469) | 5/49 (10.2) | n/a |  |  |  | 5/52 (9.6) |
| focal seizures (HP:0007359) | 8/49 (16.3) | n/a | + |  |  | 9/52 (17.3) |
| absent | 14/49 (28.6) | n/a |  | + |  | 15/52 (28.8) |
| no information available | 35/84 (41.7) | n/a |  |  |  | 35/87 (40.2) |
| **Neuroophthalmological features** | **24/27 (88.9)** | **1/5 (20.0)** | + |  | + | **27/35 (77.1)** |
| strabismus (HP:0000486) | 13/27 (48.1) | n/a | + |  |  | 14/35 (40.0) |
| nystagmus (HP:0000639) | 2/27 (7.4) | n/a |  |  | + | 3/35 (8.6) |
| strabismus and nystagmus (HP:0000486, HP:0000639) | 4/27 (14.8) | n/a |  |  |  | 4/35 (11.4) |
| optic nerve hypoplasia (HP:0008058) | 5/27 (18.5) | 1/5 (20.0) |  |  | + | 7/35 (20.0) |
| absent | 3/27 (11.1) | 4/5 (80.0) |  | + |  | 8/35 (22.9) |
| no information available | 57/84 (67.9) | 15/20 (75.0) |  |  |  | 72/107 (67.3) |
| **Facial anomalies (HP:0000271)** | **18/26 (69.2)** | **9/15 (60.0)** | + | + | + | **30/44 (68.2)** |
| micro-/retrognathia | 6/26 (23.1) | 7/15 (46.7) |  |  |  | 13/44 (30.0) |
| absent | 8/26 (30.8) | 6/15 (40.0) |  |  |  | 14/44 (31.8) |
| no information available | 58/84 (69.0) | 5/20 (25.0) |  |  |  | 63/107 (58.9) |

n/a: not applicable

We further included available likely pathogenic or pathogenic variants from ClinVar,^21^ denovo-db^22^ and DECIPHER^23^. As phenotype information was insufficient in most of these database cases, only variant information was included.

All variants were harmonized to the NM_006009.3 transcript of the GRCh37/hg19 human reference genome based on Human Genome Variation Society (HGVS) recommendations using the Mutalyzer^24^ web services. To ensure consistency in the clinical interpretation we independently applied the American College of Medical Genetics and Genomics (ACMG) criteria^25^ to all variants with the WGLAB InterVar-tool^26^.

### Protein structure analysis of the tubulin alpha-1A variants

Using R and ggplot2^27^ we analyzed spatial distribution of all variants in the linear gene model to provide an insight into the variant distribution. Utilizing Pymol (Version 1.8.6.0; Schrödinger, LLC) installed through Conda (Version 4.4.9 build 3.0.27 with Python 2.7.14; Anaconda Inc.) publicly available tertiary protein structure data of TUBA1A (PDB-ID: J5CO^28^) was used to classify variants in different groups of potential functional effects as suggested previously.^29^ This classification is based on the interaction of the tubulin monomer with neighboring tubulin proteins within the polymer (heterodimer, protofilament, microtubule), with MAPs, or motor proteins. While functional evidence was present for a minority of the variants^5,30^, most mutational effects are based on localization-dependent predictions. As a template we used 51 already classified *TUBA1A* variants^31^ likely affecting the binding of microtubule associated proteins (“MAP binding”) or motor proteins, the tubulin folding (“Tubulin folding”), heterodimer and microtubule stability (“Intradimer interaction” and “Longitudinal interaction”) the formation of the hollow tubular structure of the microtubule (“Lateral interaction”)^32,33^ or microtubule dynamics, protein folding and heterodimer stability (“GTP [Guanosintriphosphat] binding”)^29,32^. The specific detrimental effect of variants facing the luminal protein surface (“Lumen facing”) is currently unknown.

### Computational analyses of *TUBA1A* missense variant spectrum

We here analyzed the ability of six different computational classifiers (three ensemble scores: CADD, M-CAP, REVEL and the three commonly used scores Polyphen-2, SIFT, MutationTaster) to discriminate pathogenic and neutral population variants by generating all possible missense variants for *TUBA1A*. First all single base exchanges were generated in the *TUBA1A* gene region of the GRCh37/hg19 reference (chr12[hg19]:49578578-49583107) as variant call format (VCF) file. These were then annotated with computational scores and databases from dbNSFP^34^ version 2.9.3 and variant frequencies from the gnomAD database^35^ version 2.0.1 using SnpEff/SnpSift^36^. Missense variants affecting the NM_006009.3 transcript of *TUBA1A*, excluding variants, which were additionally annotated as potentially affecting splicing, were selected. R language^37^ version 3.4.3 with RStudio IDE version 1.1.383 (RStudio, Inc.) with packages from the tidyverse/ggplot2^27^ collection were used for plotting and analysis of this variant data provided in the File S1. To analyze possible mutational hotspots, we generated density plots of pathogenic missense variant frequencies reported in the literature and missense variants reported in controls from gnomAD with the “geom_density” function (“adjust” parameter set to 1/4) in ggplot2. To analyze protein regions of higher conservation we plotted all missense variants sorted by amino-acid position with each respective computational score and fitted generalized additive models using the “geom_smooth” function in ggplot2 to produce a smoothed line. Additionally, variants and scores were plotted as scatter and violin plots and two-sided Wilcoxon signed-rank test from the ggsignif package was used to determine whether there was a statistically significant difference between four different missense variant groups (“clinical review”, “database”, “gnomAD”, “simulated”). “Clinical review” included variants from individuals with available phenotype information from our literature review and the three cases reported here, “database” included (likely) pathogenic variants from databases like ClinVar without clinical information, “gnomAD” included all variants present in healthy controls without neurodevelopmental disorders from the gnomAD database, and “simulated” all other possible missense variants in *TUBA1A*.

### Analysis of genotype-phenotype relation

We used the curated set of clinical information and corresponding harmonized variant information to analyze a possible genotype-phenotype relationship by comparing the radiological and clinical features with variant characteristics. We visualized and structured the acquired categorical data into a grid plot using ggplot2 and the tidyverse^27^ package for hypothesis formation. Based on this presentation we used the vcd package^38^ to analyze the relationship between variant characteristics and clinical data of the individuals by generating mosaic or association plots. As many values in the resulting contingency tables contained values below five, we estimated p-values using a two-sided Fisher's exact test with the “simulate.p.value” setting based on 2,000 replicates in R. One-letter amino-acid nomenclature is used in the resulting plots because of space constrains.

## RESULTS

### Results of exome sequencing in 3 affected individuals

We identified three heterozygous missense variants c.518C>T, c.1307G>A, and c.641G>A in *TUBA1A*. Segregation analysis demonstrated that all variants were *de novo*. The missense variant c.518C>T, p.(Pro173Leu) identified in individual i084n and the missense variant c.641G>A, p.(Arg214His) in individual i086n, located in exon 4 of *TUBA1A*, were both previously reported either in an affected individual with autism spectrum disorder^39^ (c.518C>T, p.(Pro173Leu)) or in several affected individuals with developmental delay and complex cerebral malformations^20,40^ (c.641G>A, p.(Arg214His)). The heterozygous missense variant c.1307G>A, p.(Gly436Asp) identified in individual i085n was absent in the unaffected parents (*de novo*, sample identity confirmed), not listed in gnomAD, located in a highly conserved domain and multiple lines of computational evidence predicted a deleterious effect. Thus, we classified all variants as pathogenic (class 5) in accordance with the ACMG criteria.

### Mutational spectrum and distribution of *TUBA1A* variants

We retrieved a total of 61 distinct variants from 84 born individuals and 20 fetuses from 28 published articles in Pubmed and 59 further distinct variants from databases.^5,13,15,20,30,39-67^ Moreover we identified one novel variant c.1307G>A, p.(Gly436Asp), not reported in databases or the literature, in one of the three herein described individuals. Of these 121 distinct variants 119 were missense and two led to a premature stop codon located at the C-terminal domain and likely to escape nonsense mediated decay. Common recurrent variants were c.1205G>A p.(Arg402His), c.1204C>T p.(Arg402Cys) and c.790C>T p.(Arg264Cys) reported 11, 8 and 10 times, respectively. The Arg402 residue is the most commonly (13.3%) affected amino-acid position (Arg402His, Arg402Cys, Arg402Leu, Arg402Ser). After standardization to the ACMG criteria, 120 of the 121 distinct variants were classified as likely pathogenic or pathogenic (ACMG class 4/5) (99.2%) and one variant (c.1224C>A, p.(Tyr408*)) was classified as of unknown significance (VUS, ACMG class 3).

TUBA1A consists of the N-terminal, intermediate and C-terminal domains.^68^ Annotation of variants on the linear gene model revealed that variants were distributed all over the *TUBA1A* gene with a statistically significant clustering around the Arg402 residue in exon 4 in the C-terminal domain. This cluster correlates with high computational prediction scores for missense variants (Fig. 1A-C; Fig. S1). Variants in the linear C-terminal region predominantly affect the binding of MAPs or motor proteins. Strikingly, computational scores for the different missense variant groups (“clinical review”, “database”, “gnomAD”, “simulated”) mostly showed no significant difference (Fig. 1D; Fig. S1). After mapping of the amino acid residues on the 3D protein structure, we observed that most unique variants in “clinical review” (n=121) are predicted to compromise tubulin folding (34.7%) or possibly affecting the interaction with MAPs or motor proteins, such as kinesins and dyneins (24.8%) (Fig. 2). A minority of variants is predicted to affect longitudinal (8.3%), lateral (8.3%) and intradimer (7.4%) interactions, respectively. Finally, 14% of variants are lumen facing and only 2.5% likely affect GTP binding. Considering all assembled variants including the recurrent ones (n=166), the majority (38.6%) is predicted to impair the interaction of MAPs or motor proteins. Of these, 22 affect the Arg402 position. Variants identified in the three individuals i084n, i085n, i086n described here are predicted to affect tubulin folding (c.518C>T, p.(Pro173Leu), MAP binding (c.1307G>A, p.(Gly436Asp) and intradimer interactions (c.641G>A, p.(Arg214His), respectively.

**Figure 1.**
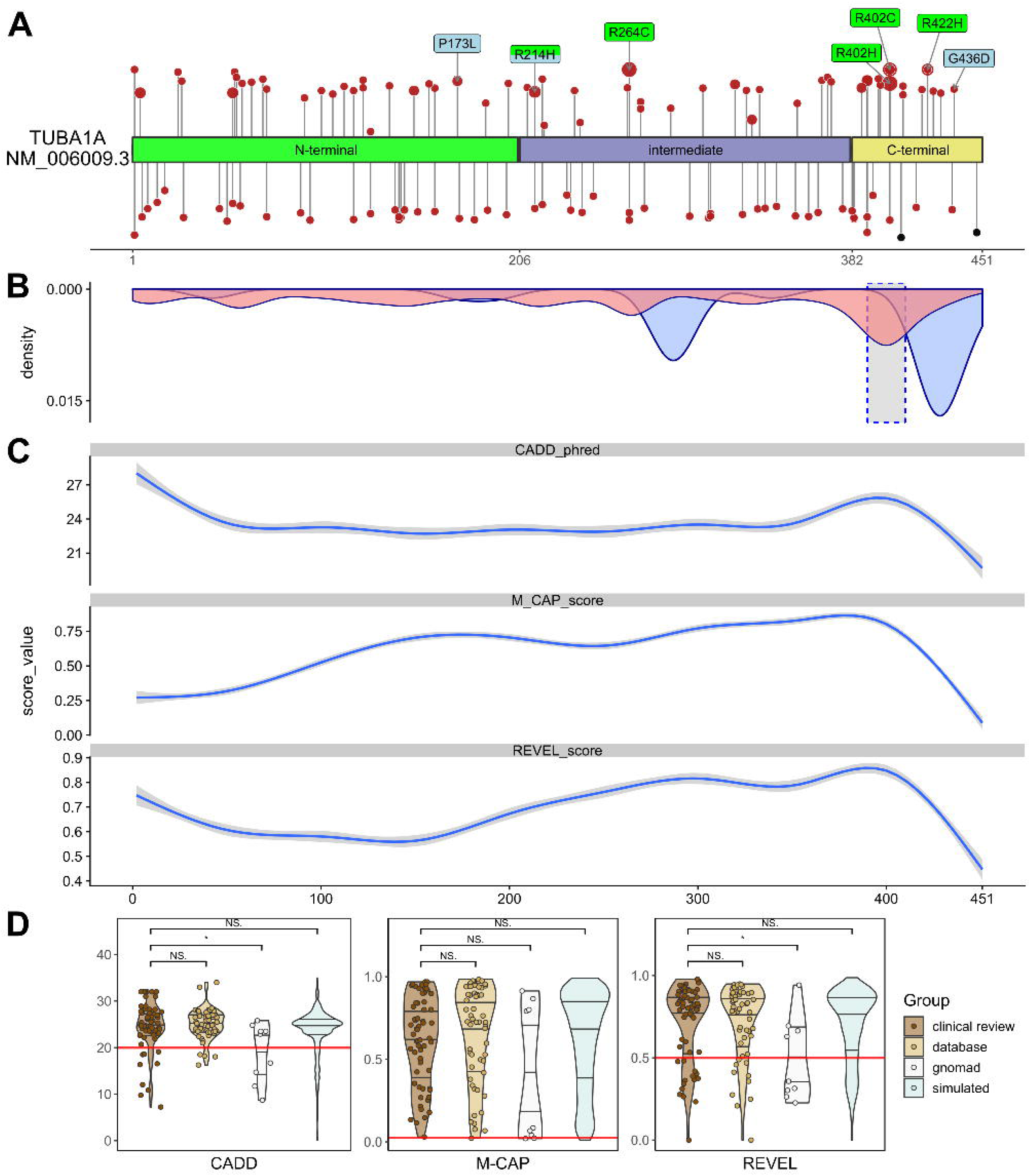
Distribution and computational scores of TUBA1A variants. (A) TUBA1A domains and localization of variants (missense variants in red, truncating variants in black). Variants above protein scheme are from published data in PubMed, below from databases (ClinVar, DECIPHER, denovo-db). Variants reported ≥ 3 times (green) and from the cases reported here (blue). While the size of the circle is proportional to the reported frequency, the height is proportional to the CADD-score. (B) Density plot of all missense variants (pathogenic in red, present in gnomAD in blue). The dashed highlighted grey box indicates the region around Arg402 with significant clustering of pathogenic variants (see Fig. S2). (C) Generalized additive models of the CADD, M-CAP and REVEL scores for all possible missense variants (see also Fig. S1 A). (D) Violin- and scatter-plot comparing the three computational scores for missense variants found in two clinical groups of individuals (“clinical review”: 104 cases from literature review and the three cases reported here for a total of 62 distinct variants; “database”: 59 individuals from ClinVar, denovo-db and DECIPHER for a total of 59 variants), healthy controls (“gnomAD”: 9 variants) and all other possible missense variants (“simulated”: 2841 variants) (see also Fig. S1 B).

**Figure 2.**
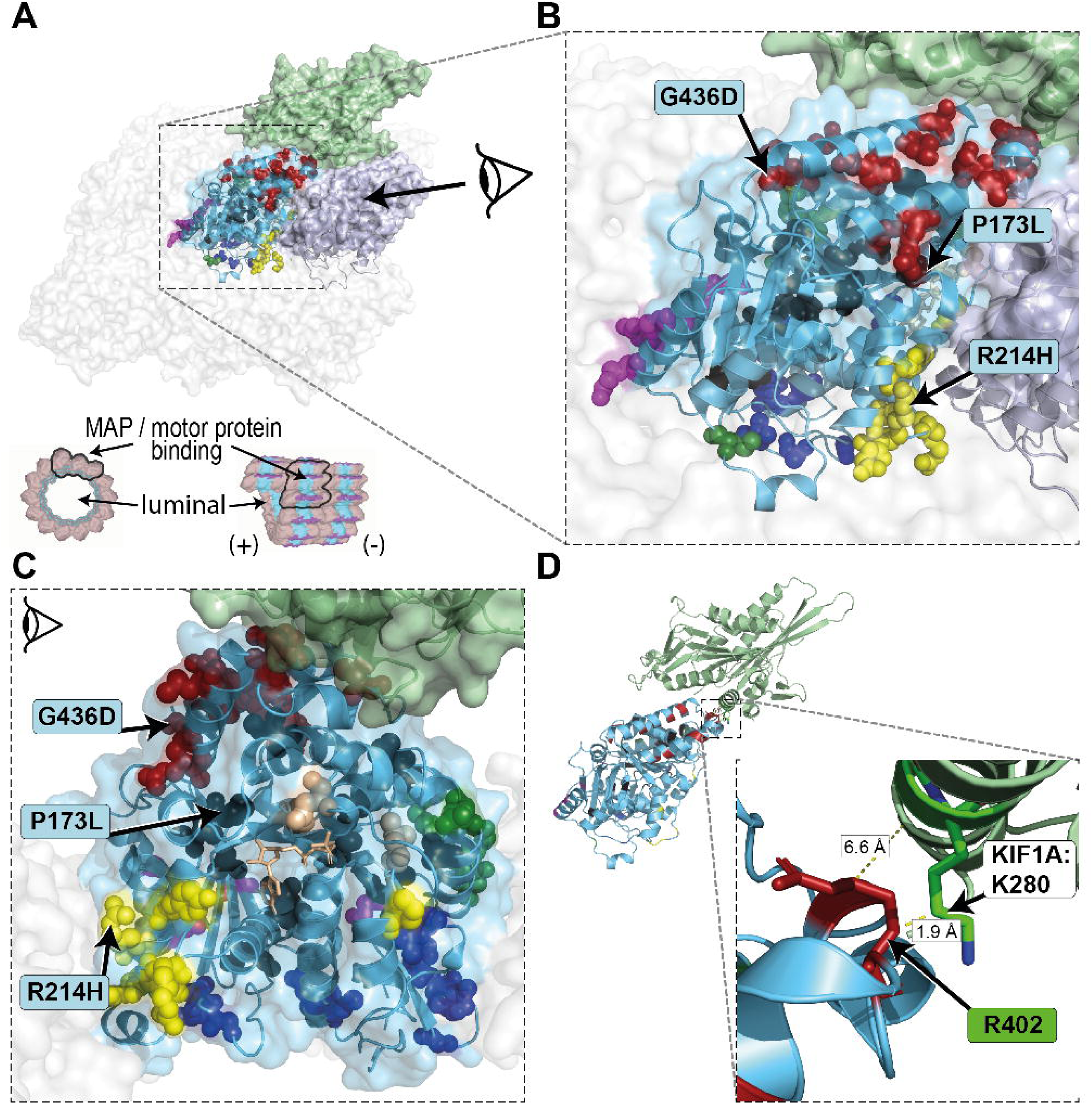
Mapping of reported variants onto 3D structure of tubulin alpha-1A. (A) TUBA1A (light blue) monomer in the center surrounded by TUBA1A monomers to the lateral sides and TUBB3 monomers to the longitudinal sides (transparent surfaces). The TUBA1A (light blue) - TUBB3 (grey) heterodimer is highlighted and shown in ribbon representation (based on PDB: 5JCO^28^). Exemplary for a motor protein KIF1A (green; PDB: 2HXF^72^) is shown interacting on the external surface. Mutated residues are shown in spheres and likely affect the binding of MAPs or motor proteins (red), tubulin folding (black), intradimer interactions (yellow), longitudinal interactions (magenta), lateral interactions (green) or GTP-binding pocket (beige). Variants on the luminal side are shown in blue. A cross section and longitudinal view of a microtubule^73^ is provided for orientation. (B) Close-up view of the central TUBA1A monomer and (C) lateral-view with TUBB3 removed from the dimer. The GTP molecule (beige), required for polymerization, is presented in stick representation. Variants identified in the three individuals i084n (P173L), i085n (G436D), i086n (R214H) described here affect tubulin folding, MAP binding and intradimer interaction, respectively. (D) Simplified representation of TUBA1A and KIF1A with protein surface and spheres removed. The amino acid residue R402 (red stick representation) of TUBA1A is localized near the KIF1A protein, in particular to the amino acid residue K280 (minimal distance 1.9 Å; green stick representation).

### Clinical spectrum of *TUBA1A* variants

Based on available information, major neuroradiological features of TUBA1A-associated tubulinopathy include anomalies of the cortical gyration (99.0%, 95/96), with lissencephaly and polymicrogyria reported in 70.0% (67/96) and 18.8% (18/96) respectively. Further anomalies affect the basal ganglia (98.3%, 58/59), the corpus callosum (96.2%, 102/106), the capsula interna (96.2%, 25/26) and the cerebellar vermis (94.0%, 78/83). Ventricular dilatation was reported in 100.0% (49/49) and anomalies of the hippocampus in 78.4% (30/38) (Table 1). Clinical features included developmental delay (98.1%, 52/53), microcephaly (76.0%, 57/75), epilepsy (71.2%, 37/52) and spasticity (62.5%, 25/40) (Table 2). Data missingness ranged from 0.9% (corpus callosum) to 75.7% (internal capsule) for neuroradiological features and from 29.9% (microcephaly) to 67.3% (neuroophthalmological features) for clinical features. We provide a detailed summary of the currently described clinical features in born individuals and fetuses with details of data missingness in Tables 1 and 2.

### Relation between Genotype and Phenotype

We used the clinical information of the 104 individuals from the “clinical review” group and the herein described three patients (total n=107) to analyze a possible relationship between genotype and phenotype. Individuals with recurrent variants, mostly affecting MAP binding, show a similar phenotype combination in the matrix plot (Fig. 3A; see also Fig. S4). Patients with the missense variant p.(Arg402Cys) are mostly described with a cortical-gyration pattern of agyria-pachgyria (“Ag-Pg”), dysplastic corpus callosum (“D”), a cerebellar vermis hypoplasia (“H”) and have no information reported for the brainstem.

**Figure 3.**
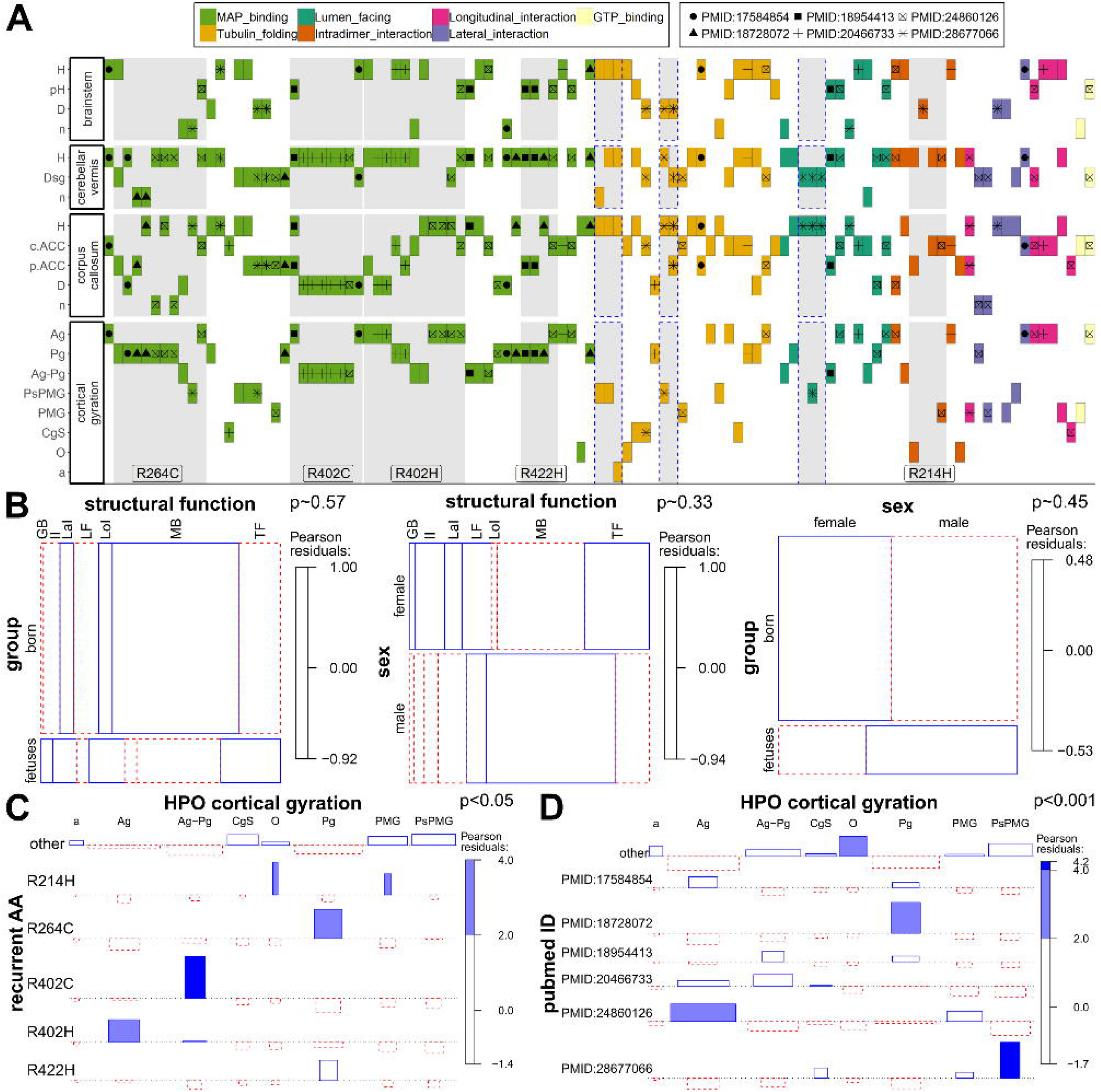
Genotype-phenotype analysis. (A) Different colors indicate the functional class of the amino acid residue in structural model (legend 1). Different symbols indicate the PubMed identifier (PMID) of publications describing ≥ 5 individuals (legend 2). Individuals described here or in the literature are sorted on the x-axis by variant functional class, localization and publication. On the y-axis phenotype categories with at least 60% data availability are presented (see also Fig. S4). Grey highlighted boxes indicate variants at the same amino acid position (also labeled) and boxes with dashed lines indicate related individuals with the same variant. (B) Mosaic plots showing the relations between individual groups (fetuses, born), variant structural function (MAP_binding = “MB”, Tubulin_folding = “TF”, Lumen_facing = “LF”, Intradimer_interaction = “II”, Longitudinal_interaction = “LoI”, Lateral_interaction = “LaI”, GTP_binding = “GB”) and sex of the individual (female = f, male = m). (C) Association plot showing the relation between recurrently affected amino-acid positions (recurrent_AA) and the neuroradiological feature of cortical gyration (pachygyria = “Pg”, polymicrogyria = “PMG”, perisylvian polymicrogyria = “PsPMG”, cortical gyral simplification = “CgS”, agyria = “Ag”, other = “O”, absent = “a”). This example (see Fig. S5) indicates a possible genotype-phenotype correlation for certain recurrent variants. (D) Association plot showing the relation between publications describing ≥ 5 individuals (“pubmed_ID”) and the neuroradiological feature of cortical gyration. This example (see also Fig. S6) indicates a probable reporting bias for this clinical feature. Two-sided Fisher's exact test has been used to estimate the presented p-values.

Because prenatally diagnosed fetal cases show a more severe phenotype than born individuals, we analyzed a possible contribution of variant characteristics to this observation. The missense variants reported in fetuses and in born individuals showed no significant difference in structural classification (Fig. 3B) and the computational scores did not significantly differ in these two groups (Fig. S3). In addition, the structural groups of missense variants were not overrepresented in females or males and the gender was also not associated with prenatal diagnosis (Fig. 3B).

The visual inspection of the matrix plot (Fig. 3A; Fig. S4) indicated that certain clinical features are enriched in individuals with recurrent variants. Indeed, explorative comparison between missense variants at recurrent and non-recurrent positions confirmed differences in reported clinical features of the individuals carrying these missense variants (Fig. 3C; Fig. S5). Despite our effort to collect all variants and clinical information described for *TUBA1A-* associated tubulinopathy, we did not obtain enough data to further analyze the phenotype differences for these variants.

Finally, we observed that individuals with the same variants and similar phenotypes were often reported together (e.g. Fig. 3A “+” symbol for Arg402Cys reported 5 times in PMID:20466733^30^). Regarding this observation, we found a significant difference in the use of clinical descriptions in publications describing multiple individuals. Kumar et al. (PMID:20466733^30^) and Bahi-Buisson et al. (PMID:24860126^13^) both describe four cases with the *de novo* missense variant c.1205G>A p.(Arg402His), but Bahi-Buisson et al. more often describes agyria (“Ag”) as cortical gyration pattern. Romaniello et al. (PMID:28677066^57^) describe perisylvian polymicrogyria (“PsMPG”) as cortical gyration pattern for four (all with different missense variants) of their 14 reported individuals’ variants while this term is only used for 6 other individuals in the entire “clinical review” group (Fig. 3D; Fig. S6).

## DISCUSSION

In this study, we identified three *de novo* missense variants in *TUBA1A* in three individuals with developmental delay and brain malformations. Since the first identification of disease-causing variants in *TUBA1A* in 2007 in two affected individuals with cortical dysgenesis,^5^ at least 121 distinct heterozygous variants in a total of at least 166 patients including our 3 affected individuals are now described. Our efforts to systematically reanalyze published data enabled insights into the current state of information about TUBA1A-associated tubulinopathy.

Anomalies of the corpus callosum ranging from partial to complete agenesis or hypoplasia are with 96.2% (102/106) the predominantly reported feature of TUBA1A-associated tubulinopathy. Cortical anomalies are the second leading clinical feature reported in 95/96 individuals (99.0%) followed by dysgenesis of the basal ganglia in 58/59 (98.3%). Two of the newly described individuals also presented these features, while individual i086n additionally manifested unilateral optic nerve hypoplasia, a feature predominantly linked to *TUBA8-* associated tubulinopathy^6^ but also present in 7/35 (20.0%) of individuals with *TUBA1A-* associated tubulinopathy.

Analysis of the type and localization of all possible 2,969 missense variants from the simulation showed that the large majority of *TUBA1A* missense variants are predicted to be deleterious (CADD ≥ 20: 84.2%, M-CAP ≥ 0,025: 98.0%, REVEL ≥ 0,5: 78.8%; Fig. 1D). This is in agreement with an ExAC Z-score^35^ of 6.23, confirming that *TUBA1A* is extremely depleted of missense variants in the general population. This resulted in high computational prediction scores independent of causality. Thus, variants might be reported to be pathogenic (ACMG class 5) despite relatively low computational scores and variants found in healthy controls might have scores above the recommended respective thresholds (Fig. 1D). After analyzing the relation of three ensemble computational prediction scores and expected pathogenicity, we concluded that computational prediction scores are of limited utility for predicting pathogenicity in *TUBA1A*. We suggest that segregation with the disease in the family or *de novo* occurrence, two major criteria of the ACMG guidelines for variant interpretation, are more appropriate for variant classification.

Based on the observation of the mutational distribution we analyzed a possible relationship between genotype and phenotype. We observed clustering of disease causing variants in the region around the amino acid residue Arg402 (Fig. 1A-C, Fig S2). The residue Arg402 is located in the interaction site of various MAPs or motor proteins^30^ which are involved in different processes including the polymerization and stabilization of microtubules and intracellular vesicle transport.^69^ Defects in some MAPs or motor proteins result in a similar clinical spectrum as observed for specific MAP-associated *TUBA1A* variants.^30,70^ Overall, variants of the Arg402 residue and other specific recurrent variants, which are predominantly MAP interacting, were previously associated with overlapping neuro-radiological features.^13,30^ Indeed, we could show a non-uniform distribution for reported clinical features and the recurrent variants (Fig. 3C), indicating a possible genotype-phenotype relation. This observation might in part be attributed to detailed structured morphological categorization of brain anomalies used by different authors and individual preferences for certain terms. In addition, difficulties in the interpretation of the radiographic cortical and subcortical anomalies or technical differences in brain imaging could represent a possible confounder. Of note, recurrent variants with similar phenotype combinations were often reported by the same authors indicating a possible observational bias (Fig. 3D), thus limiting the interpretation of these genotype-phenotype relations. Another problem hindering a more detailed investigation is the high degree of missing data we recognized for several phenotypic categories. The directed acyclic graphs structure of HPO allows grouping of specialized terms into less specialized parent terms. Future development of algorithms comparing the phenotypic similarity between groups of individuals with the same or functionally similar pathogenic variants might alleviate some of these problems and allow further characterization of variant specific phenotypes. However, some of these endeavors could be hampered by the difficulty to distinguish between missing information and normal phenotype in published reports. This is especially problematic as HPO describes “phenotype abnormalities” but has no terms for normal phenotypes. We propose standardization in clinical reporting of rare disease cases based on expert recommendations with a minimal scheme covering disease specific phenotypes.

Even though *TUBA1A-associated* tubulinopathy is the most common tubulinopathy form, our results indicate that more clinical and mutational information is necessary to evaluate a potential genotype-phenotype correlation. This became apparent in fetuses, where we and others observed the most severe phenotypic spectrum compared to born cases.^13,20^ This could not be explained by specific properties of the identified variants (Fig. 3B, Fig. S3). We therefore propose that additional variants in other genes or random developmental processes in cellular pathways in the respective individuals are underlying the phenotypic variability. Genome wide and functional studies might help to allow further characterization into specific clinical groups.

## CONCLUSION

Our systematic reanalysis of published clinical data allowed an explorative investigation of a genotype-phenotype relationship. We found an enrichment of specific radiological features in recurrent variants; however, insufficient data availability, data variability and a possible observer bias were limiting factors for possible associations. A thoroughly conducted clinical examination and the standardized reporting of phenotype and genotype information in online databases, e.g. ClinVar^21^ and LOVD^71^ are fundamental for the systematic analysis of rare diseases such as *TUBA1A-associated* tubulinopathy.

## ACKNOWLEDGEMENTS

We thank all participants and their families for taking part in this study. We acknowledge the excellent technical support of Angelika Diem, and Heike Friebel-Stange. We also thank the Exome Aggregation Consortium and the groups that provided exome variant data for comparison. A full list of contributing groups can be found at http://exac.broadinstitute.org/about. This study utilized data generated by the DECIPHER community. A full list of centers that contributed to data generation is available online from http://decipher.sanger.ac.uk and via email from.

This study was supported by grants from the German Research Foundation (DFG; grants TH 896/3-4), the Interdisciplinary Centre for Clinical Research (IZKF) of the Friedrich-Alexander-Universität Erlangen-Nürnberg (Project F4) and by the ELAN Fonds (14-08-06-1) of the Faculty of Medicine of the Friedrich-Alexander Universität Erlangen-Nürnberg (FAU) to CT.

**ABBREVIATIONS**
ACMG: : American College of Medical Genetics and Genomics
CMA: : Chromosomal microarray
GTP: : Guanosintriphosphat
HGVS: : Human Genome Variation Society
HPO: : Human Phenotype Ontology
MAPs: : Microtubule-associated proteins
MRI: : Magnetic resonance imaging
OFC: : Occipitofrontal circumference
VCF: : Variant call format

## AUTHORS’ CONTRIBUTIONS

CT and UH provided all clinical findings and patient samples. MK conducted Array analyses. CK, SU, AE, CT, AR and BP analyzed and interpreted the sequencing data. BP created Fig. 1+3 and the supplementary materials. MH performed the literature review and standardization to HPO terms. MH and BP created Fig. 2. BP, MH and CT wrote and edited the manuscript, CK, MK, SU, AE, CT, AR and BP reviewed the draft manuscript.

## AVAILABILITY OF DATA AND MATERIALS

All data generated or analysed during this study are included in this published article [and its supplementary information files].

## WEB RESOURCES

gnomAD browser: http://gnomad.broadinstitute.org/ Mutalyzer: https://mutalyzer.nl/ wInterVar: http://wintervar.wglab.org/

